# Development of amyloid beta gold nanorod aggregates as optoacoustic probes

**DOI:** 10.1101/2021.10.25.465704

**Authors:** Mahmoud G Soliman, Hannah A Davies, Jack Sharkey, Raphaël Lévy, Jillian Madine

## Abstract

Propagation of small amyloid beta (Aβ) aggregates (or seeds) has been suggested as a potential mechanism of Alzheimer’s disease progression. Monitoring the propagation of Aβ seeds in an organism would enable testing of this hypothesis and, if confirmed, provide mechanistic insights. This requires a contrast agent for long-term tracking of the seeds. Gold nanorods combine several attractive features for this challenging task, in particular, their strong absorbance in the infrared (enabling optoacoustic imaging) and the availability of several established protocols for surface functionalization. In this work, polymer-coated gold nanorods were conjugated with anti-Aβ antibodies and specifically attached to pre-formed Aβ seeds. The resulting complexes were characterized for their optical properties by UV/Vis spectroscopy and multispectral optoacoustic tomography. The complexes retained their biophysical properties, i.e. their ability to seed Aβ fibril formation. They remained stable in biological media for at least 2 days and showed no toxicity to SH-SY5Y neuroblastoma cells up to 1.5 nM and 6 μM of gold nanorods and Aβ seeds, respectively. Taken together, this study describes the first steps in the development of probes for monitoring the spread of Aβ seeds in animal models.

## Introduction

Aggregation and deposition of Aβ and tau proteins occurs in the brain in Alzheimer’s disease and is linked to neurodegeneration [1]. However, the trigger for Aβ and tau deposition remains unknown. Recent research has highlighted pathological similarities between amyloid and prion diseases [2-4]. In prion disease, the mis-folded protein acts as a template or ‘seed’ to convert the native protein to a mis-folded pathogenic form. Growing evidence suggests that a similar ‘seeding’ mechanism could be associated with additional amyloid disorders such as Alzheimer’s disease [3], Parkinson’s disease [5], Huntington’s disease [6], AA amyloidosis [7] and ATTR amyloidosis [8].

The body of evidence supporting Aβ having prion-like properties and Aβ-associated diseases spreading via seeding is rapidly expanding. Early work showed that injection of Alzheimer’s disease brain homogenates into young APP23 mice overexpressing amyloid precursor protein (APP) results in accelerated disease pathology observed by the presence of Aβ plaques at an earlier age than control animals [4, 9]. Whilst in the initial work, seeds were injected directly into the brain, a further study has shown that peripheral injection of seeds into the peritoneal cavity can also accelerate disease onset [10]. Support for a similar seeding mechanism in human patients was reported from post-mortem analysis of a cohort of patients that had received growth hormone injections, derived from cadavers, as children. A sub-set of this cohort developed Creuzt-Feldt Jacob disease (CJD), a well-known prion disease, as a result of growth hormone contamination with prion seeds. In 2015, Jaunmuktane, et al., reported that 4/8 patients also had signs of Aβ deposition consistent with early onset of Alzheimer’s disease and suggested that these patients could have contracted Alzheimer’s disease in a similar mechanism to CJD, i.e. a prion-like infection with Alzheimer’s disease seeds from cadaveric tissue [11], although those results are debated [12-14].

Evidence for the transmission of amyloid proteins is not confined to neurodegenerative diseases. Several groups have demonstrated the transmissibility of AA amyloidosis through a variety of administration routes and across species [15-17]. Furthermore there are suggestions that seeding mechanisms play a role in the development and progression of ATTR amyloidosis which affect treatment and prognosis [8]. In all these diseases and examples, a critical currently unknown factor is how seed species trigger disease, specifically how seed species injected at one site (e.g. peritoneal cavity of mice) can affect amyloid deposition elsewhere in the body (e.g. brain). To investigate this puzzling phenomenon, novel experimental approaches to track seed species from site of injection to site of action are needed. This would result in enhanced understanding of the mechanisms that underpin disease initiation and progression. As a first step towards this ambitious goal, we present in this article the preparation of seeds labelled with gold nanorods for future tracking with multispectral optoacoustic tomography (MSOT).

MSOT is a real-time optical imaging technique that provides a non-ionizing and noninvasive imaging modality which relies on the photoacoustic effect, which was first observed by Alexander G Bell in 1880 [18]. MSOT has been used for several preclinical and clinical applications [19-21] because of its high spatial resolution (∼150 μm) and penetration depth (∼ 5 cm)[22]. In optoacoustic tomography, a short near infrared laser pulse is partially absorbed by endogenous molecules or exogenous contrast agents causing a rapid thermoelastic expansion, which generates ultrasound waves. These are detected by ultrasonic transducers placed outside the tissue [23]. MSOT relies on a multiwavelength excitation and subsequent spectral deconvolution to identify endogenous molecules or exogenous contrast agents [22]. Therefore, agents with strong and distinctive absorption spectra in the near infrared are best suited for MSOT imaging.

Gold nanoparticles have strong optical absorption that results from the surface plasmon resonance. The position of the absorption band can be tuned through changes in nanoparticle size and shape. Anisotropic nanoparticles, e.g., gold nanorods, exhibit two bands, the transverse plasmon band located in the visible region around 520 nm, and the longitudinal band located in the near infra-red region, with the exact wavelength tunable by controlling the aspect ratio of the nanorods [24]. For this reason, gold nanorods have been successfully used as contrast agents for MSOT [25, 26]. In this work, we cross-link anti-Aβ antibodies to gold nanorods and demonstrate that those conjugates selectively associate with Aβ seeds. The resulting seeds-Abs-GNR complexes can serve as optical probes for MSOT imaging to monitor the propagation of Aβ seeds in *in vivo* murine models.

## Materials and Methods

### Materials

Prior to use, all glassware was washed with aqua regia and rinsed thoroughly with milli-Q water. All chemicals were used as received. Hydrogen tetrachloroaurate (III) hydrate (HAuCl_4_), hexadecyltrimethylammonium bromide (CTAB), sodium borohydride (NaBH_4_), ascorbic acid, Ethyl-3-(3-dimethylaminopropyl)carbodiimide (EDC)/N-hydroxysulfo-succinimide (sulfo-NHS), 2-(N-morpholino)ethanesulfonic acid (MES), poly(isobutylene-maleic-altanhydride), hydrochloridric acid, 5-bromosalicylic acid, silver nitrate (AgNO_3_), phosphine buffered saline (PBS), Ethylenediaminetetraacetic acid (EDTA), resazurin, and dodecylamine, α-metoxy-ω-thiol-poly(ethylene glycol) (mPEG-SH (PEG_2k_), Mw = 2 kDa) HAMS-F12, Non-essential amino acids (NEAA), Penicillin-Streptomycin (Pen-Strep), mouse serum and Foetal Bovine Serum (FBS) were purchased from Sigma-Aldrich (UK). All cell culture plasticware was purchased from Corning Inc. (Corning, NY, USA).

### Synthesis of gold nanorods

Gold nanorods with core diameter and length of ca.15 and 60 nm, respectively, were prepared by the seed-mediated growth method by two steps (seed and growth solutions) as reported previously [27] Briefly, seed solution of less than 4 nm gold nanoparticles was prepared by mixing 5 mL of 0.5 mMHAuCl_4_ with 5 mL of 0.2 M CTAB in a glass vial. To this solution, 0.6 mL of freshly prepared NaBH_4_ diluted to 1 mL with deionized water was added with strong stirring for 2 min. The resulted brown-yellow color solution was kept at room temperature for 15 min before use. To prepare a growth solution, 9.0 g of CTAB and 1.1 g of 5-bromosalicylic acid were dissolved in 250 mL of warm water in a 500 mL Erlenmeyer flask. After that, the solution was cooled down to 30 °C, and 18 mL of 4 mMAgNO_3_was added. The mixture was kept at 30 °C for 15 min and then, 250 mL of 1 mM HAuCl_4_ solution was added. After 15 min of stirring (400 rpm), 2 mL of 0.064 M of ascorbic acid was added, and the solution was vigorously stirred for 30 s until it became colourless. Finally, 0.4 mL of freshly prepared seed solution (as described above) was injected into the growth solution and stirred for 30 s. Finally, the solution was kept overnight at 30 °C without stirring to allow the gold nanorods to grow. Next day, the resulted gold nanorods were subjected to one-time cleaning by centrifugation (8960xg, 30 min). Thereafter, the supernatant was discarded and the pellet was dispersed in deionized water and stored for further use.

### Phase transfer and polymer coating

For polymer coating of gold nanorods, the aqueous solution was first subjected to a phase transfer from aqueous medium to organic solvent following the protocol previously reported [28] Briefly, a specific amount of PEG_2k_ in water (see Table S1) was injected into the solution of gold nanorods and further stirred at room temperature for 24 h. After that, a 0.4 M solution of dodecylamine in chloroform was added and the mixture stirred at 1200 rpm until the gold nanorods was transferred to the chloroform phase. For purification, the particles were centrifuged twice (see Table S1 for parameters) to remove unbound/unreacted dodecylamine and PEG_2k_. Then, the gold nanorods were dispersed in chloroform and stored for further use. Their concentration was calculated using Beer Lambert law (A_450_ = c_NP_ ε(450 nm) l_path_) with UV/Vis absorbance determined at wavelength of 450 nm (A_450_) and extinction coefficient (ε(450 nm) = 2.11·10^9^[M^-1^cm^-1^]) as reported for gold nanorods of very similar dimensions [29]. Measurements were carried out with a cuvette path length of l_path_ = 1 cm.

The dodecylamine-capped gold nanorods were transferred from chloroform to water by applying polymer coating technique using dodecyl–grafted-poly(isobutylene-alt-maleic anhydride)(in the following referred to as PMA) as described by Lin et al.[30]. For details about the synthesis of the amphiphilic polymer used here we refer to previous work [30]. The amount of added PMA per NP was calculated as described in the Supporting Information (appendix). The polymer coating was then carried out according to the protocol previously described [28]. A schematic illustration describes the steps of NPs’ surface modification (Scheme 1).

**Scheme 1:**
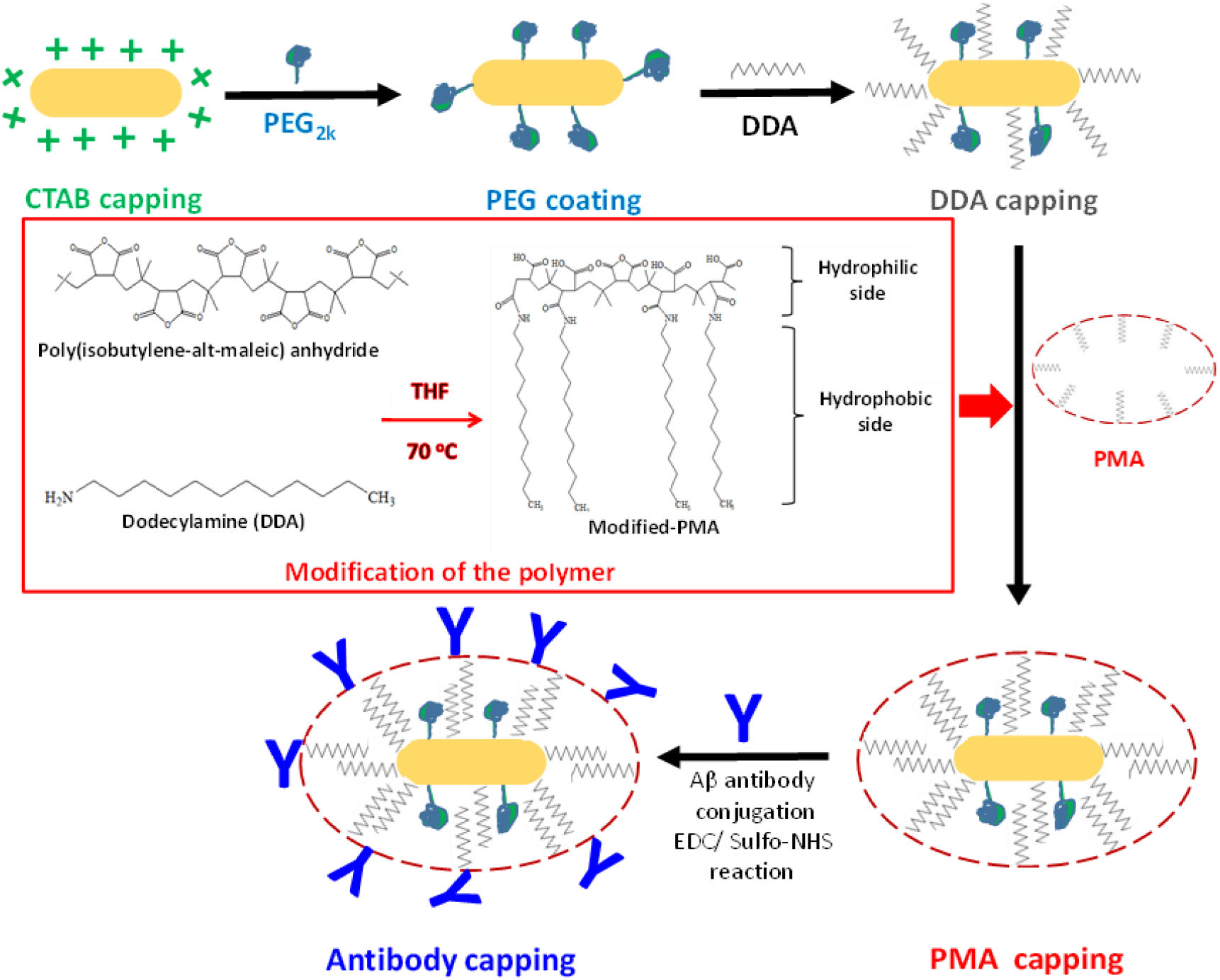
Schematic illustration shows the steps of NPs’ surface modification.

### Gold nanorods functionalization with antibodies

After the polymer coating with PMA, the gold nanorods’ surface is rich in carboxylic groups (–COOH) that can be linked to Aβ antibodies (Abs-Aβ) using EDC/sulfo-NHS coupling reaction (Scheme 1)[31]. To achieve high conjugation efficiency, the concentrations of coupling reagents (EDC/Sulfo-NHS) and Abs-Aβ were optimized sequentially. The molar ratio between EDC and sulfo-NHS was fixed at 1:2, and the amount of Abs-Aβ added to the reaction was introduced with the same volume as described in the following coupling reaction. Briefly, 10 μL of Abs-Aβ (5-75 μg/mL, 6E10 BioLegend) was added to a suspension of gold nanorods at a concentration of 5 nM in 10 mM MES buffer at pH 6. To this mixture, an equal volume of a solution containing 0.5-4 mmoles of sulfo-NHS and 1-8 mmoles of EDC was added (see Table S2 for more details). The reaction mixture was left at room temperature for 2 h and then was incubated for extra 6 h at 4 °C.

### Association of gold nanorods with Aβ40

Aβ40 (BioLegend) was incubated in PBS at 200 μM with agitation for 3 days at 37 °C. To produce seeds, Aβ fibrils were subjected to three cycles of ultrasonication pulses (amplitude of 10 %) for a period of 30 sec/cycle. Aβ fibrils/seeds were diluted to required concentrations (0.1-10 μM equivalent monomer concentration) and incubated with gold nanorods (5-500 pM) overnight at room temperature before using. The resulted complex (Aβ fibrils or Aβ seeds-GNRs) were characterized by UV/Vis spectroscopy and transmission electron microscope (TEM).

### Physicochemical characterization

Absorption spectra of gold nanorods with different surface coatings were acquired across the wavelength range of 400–1000 nm with UV/Vis spectrometer (Cary Eclipse).

A Malvern zetasizer Nano ZS was used to measure the hydrodynamic diameter (d_h_) and zeta potential (ζ-potential) of the gold nanorods in water with dynamic light scattering and laser Doppler anemometry respectively. The latter was also used to characterize the ζ-potentials of the prepared Aβ-seeds. All samples were equilibrated for 5 min at 25 °C to ensure motion was due to Brownian motion and not due to any thermal gradients. The data was acquired at 173° backscatter settings, using a 633 nm laser. The determined d_h_ [nm] and ζ-potentials [mV] are summarized in Table S3. Each reported value was the average of at least three independent measurements.

Gold nanorods were also characterized at each step of surface modification using a CPS disc centrifuge DC24000 (CPS Instruments Inc.). For this purpose, a gradient fluid, 8-24 wt% sucrose solution in Milli-Q water, was freshly prepared and injected in consecutive steps into the disc, rotating at a speed of 22000 rpm. Calibration was performed using poly(vinyl chloride) particles (PVC, 0.377 μm, Analytik Ltd.) as calibration standard before each measurement. Following PVC measurement, 100 μL of gold nanorods solution was injected and analyzed three times to verify data reproducibility.

The dimensions of gold nanorods, their surface modification and their interaction with Aβ species were investigated by TEM (Tecnai G3 spirit). The samples were prepared by deposition of a drop (5 μL) of diluted solution on a carbon coated copper grid and left to dry at room temperature. For negative staining samples, a drop of gold nanorods solution (incubated with Aβ-fibrils/seeds) was placed on a carbon-coated copper grid for 2 min. Following blotting, 5 μL of 4 % uranyl acetate was added for 30 sec. Size distributions were determined using ImageJ. At least 100 particles were considered to determine the average size of the different cores.

Gel electrophoresis experiments were conducted using 1% agarose gel prepared in tris-EDTA buffer at pH 7.4. Gold nanorod solutions were mixed with a specific volume of a negatively charged orange G as a gel-loading buffer to increase the density of the sample, following the protocol reported by Soliman et al. [28] and then loaded in the gel wells. Samples were run under an electric field of 100 V in tris-EDTA buffer at room temperature for 1 h using a Bio-Rad horizontal electrophoresis system.

### Gold nanorods effect on protein aggregation

Thioflavin T (ThT) fluorescence assay was used to assess fibril formation using a FlexStation3 Multi-Mode Microplate Reader (Molecular Devices). 10 μM Aβ40 with 2 μM of ThT was incubated in PBS at 37°C in a 96 well clear bottom/black wall 96 well plate. The measurements were made at regular intervals (every 5 minutes with shaking for 5 s before each read) with excitation and emission at 440 nm and 480 nm, respectively. To stop the depletion of Aβ40 from the solution resulting from non-specific binding, the plates were pre-coated with polyethylene glycol [32]. Briefly, 300 μL of 0.01 M polyethylene glycol (Mw= 300 g/mol) was added into every well of the 96-well plates and incubated at room temperature for 60 minutes. The wells were then aspirated completely and rinsed with 10 times their volume (3 ml) of Millipore ultra-pure water. The plates were allowed to dry at room temperature before use.

### Cell viability

Cell viability was tested against SH-SY5Y neuroblastoma cells at cell density of 10,000 cells/well in a 96 well plate. The cells were incubated with the particles/Aβ proteins for 24 h and their viability was evaluated by a standard resazurin assay [33]. Resazurin is a blue, non-fluorescent sodium salt, which is converted to Resorufin by metabolically active cells. Fluorescence spectra were recorded using FlexStation3 Multi-Mode Microplate Reader with excitation wavelength of 560 nm. In all tested particles/proteins, the molarity and mass concentrations are indicated. Gold nanorods molar and mass concentrations were determined based on the UV/Vis absorbance values as described in supporting information (Appendix). Viability data are presented as percentage compared to untreated cells. Data was analysed statistically using analysis of variance and Tukey’s post hoc test with p<0.05.

### Optoacoustic imaging

MSOT imaging was performed in the MSOT inVision 256-TF small animal imaging system (iThera Medical, Munich, Germany). For this purpose, an agar phantom was prepared by mixing 4 g of agar in 100 mL of hot water. Thereafter, the solution was placed in 50 mL tubes and stored at 4 °C for further use. To prepare, gold nanorods containing phantoms, different concentrations of 100 μL of gold nanorods solution was placed in phantoms wells and five cross sections of the phantom were analyzed using 15 different wavelengths in the range from 660 to 950 nm. The phantom was scanned in 2 mm steps and the following wavelengths used for acquisition; 680, 720, 760, 800, 810, 820, 830, 840, 850, 860, 870, 880, 890, 900, 950 nm. A linear based reconstruction method was applied using ViewMSOT software (iThera Medical, Munich, Germany). For multispectral images, the signal for the GNRs was unmixed by multispectral processing using the absorbance spectra for the GNRs

## Results and discussion

### Gold nanorods synthesis

Gold nanorods were prepared by a well-established procedure that results in particles capped with the toxic cationic surfactant CTAB [27]. To remove the bound CTAB, the gold nanorods were transferred from water to chloroform using both PEG_2k_ and dodecylamine. Gold nanorods exhibit characteristic absorbance peaks in both the visible and near IR regions. The positions of those peaks are sensitive to the size end geometry of the particle, but also to its local environment refractive index. Thus, a 40 nm red shift of the longitudinal plasmon band was observed (from 830 nm to 870 nm) due to the change in the refractive index of the dispersion medium from water to chloroform as shown in Fig 1a. Coating with an amphiphilic polymer (PMA-GNRs) was then used to transfer the particles back from chloroform to water and to serve as a platform for immobilization of antibodies using water-soluble cross-linkers EDC and sulfo-NHS [30]. The spectrum after transfer is similar to the starting material with a small blue shift of the peak. A range of experimental conditions were tested to maximize coupling yield estimated by the red shift of the plasmon band (Table S2 and Fig S1); in the optimized conditions a ∼5 nm red shift is observed (Fig 1b, inset)[34]. In the absence of EDC/sulfo-NHS no change in localized surface plasmon resonance (LSPR) was detected consistent with no binding occurring (Fig S1a). Electron microscopy and dynamic light scattering confirm that the particles have a narrow size distribution that is preserved during the various modification steps, with a final increase of hydrodynamic diameter upon coupling of the Aβ antibody (Fig S2 and S3, Table S3). Consistent with previous work [35, 36], the conjugation results in a reduction in the apparent diameter measured by CPS due to reduction in overall density upon antibody binding (Fig S4). Coupling of the Aβ antibody to the PMA-coated gold nanorods was further confirmed by reduced electrophoretic mobility (Fig 1c) and changes in zeta potential (Fig S5, Table S3).

**Fig 1.**
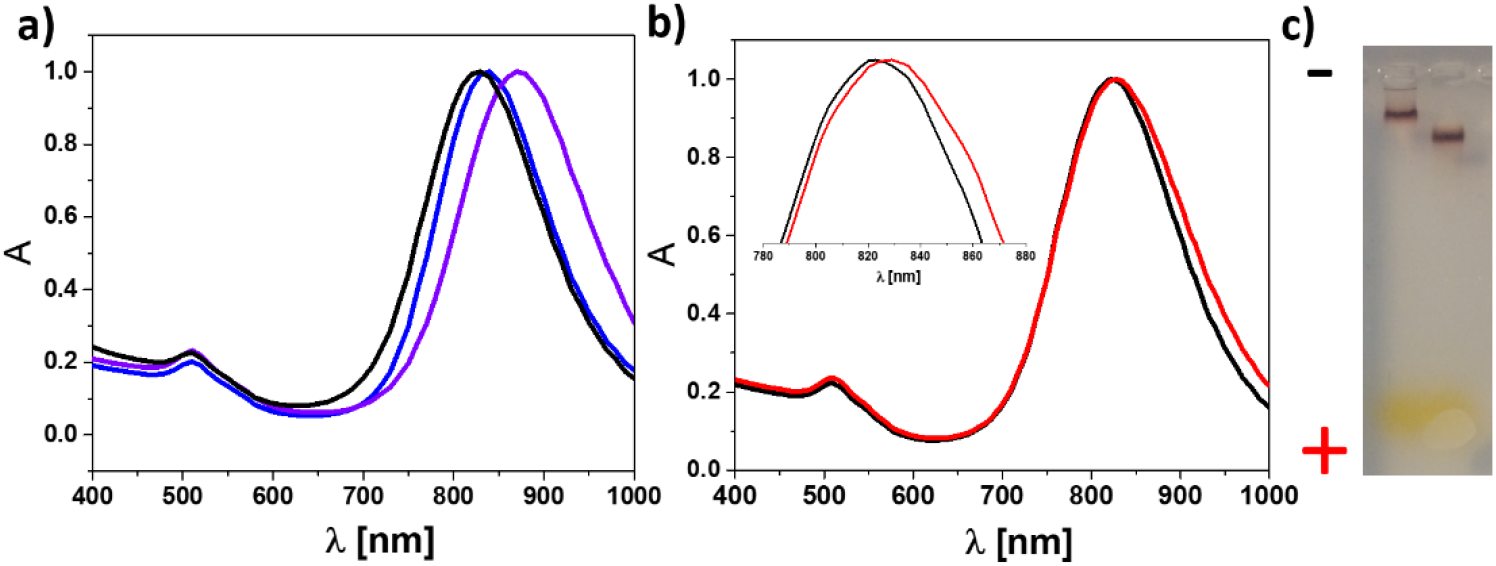
Conjugation of Aβ antibody to gold nanorods. a) Normalized absorption spectra for CTAB-capped gold nanorods (in water, blue), dodecylamine-capped gold nanorods (in chloroform, violet) and PMA-coated gold nanorods (in water, black). b) Normalized absorption spectra for gold nanorods after PMA coating (black) and after conjugation with Aβ antibody (red). Inset shows normalized data for the LSPR shift of the gold nanorods after conjugation with antibody. Gel electrophoresis of gold nanorods before (right lane, PMA-GNRs) and after (left lane, Abs-GNRs) conjugation with Aβ antibody.

### Abs-GNRs interact specifically with Aβ-40

The interaction of PMA-GNRs and Abs-GNRs with Aβ was investigated by analysing changes in the gold nanorods absorbance spectra in the presence of Aβ seeds and fibrils. No changes in the longitudinal plasmon band were observed when PMA-GNRs were incubated with Aβ seeds or fibrils (Fig 2a, 2c). In contrast, with Abs-GNRs the spectra indicated a progressive reduction in intensity indicative of association with the seeds and fibrils formation (Fig 2b, 2d). As the Fig insets show, the position of the plasmon band maxima do not shift significantly. This absence of plasmon coupling indicates that the gold nanorods remain separated from each other within the fibrillar networks [37]. This may be explained by two main factors: 1) the large steric barrier between the nanorod cores provided by the PMA and antibody layers; 2) the nanorods do not bind directly to each other but instead to another extended object, i.e. the seeds or fibrils. The preserved shape of the spectrum is an important advantage for future MSOT detection as this technique relies on multispectral unmixing to distinguish contrast agents from endogenous absorbers. Specific association of Abs-GNRs with Aβ40 is confirmed by TEM where Abs-GNRs are almost entirely co-localized with Aβ seeds and fibrils, whereas under the same conditions, a large proportion of PMA-GNRs are observed randomly scattered over the grid (Fig 3). In the representative images shown in Fig 3, ∼95% of Abs-GNRs colocalize with seeds compared with only ∼18% of PMA-GNRs. Additional images are shown in Fig S5. Incubation of Abs-GNRS overnight with seeds from a different amyloidogenic protein medin confirmed that the association between Aβ-GNRs and Aβ is specific (Fig S7).

**Fig 2.**
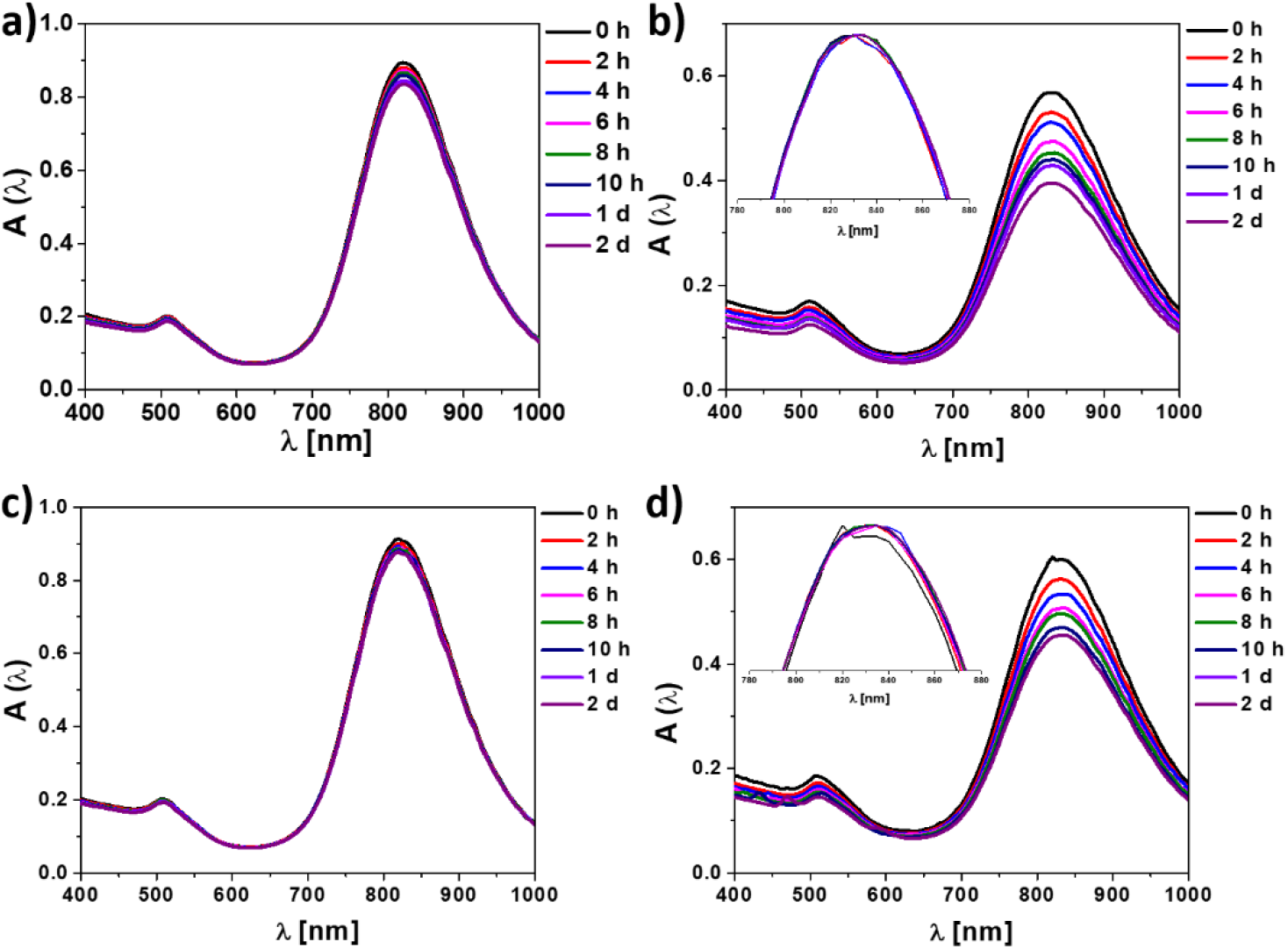
Abs-GNR show specific association with Aβ-40 fibrils and seeds. Absorption spectra (not normalized) of a) PMA-GNRs with Aβ-seeds, b) Abs-GNRs with Aβ-seeds, c) PMA-GNRs with Aβ fibrils, and d) Abs-GNRs with Aβ fibrils in water at different time-points as shown. Loss of intensity is observed due to conjugation with insoluble Aβ seeds and fibrils. Insets show normalized spectra.

**Fig 3.**
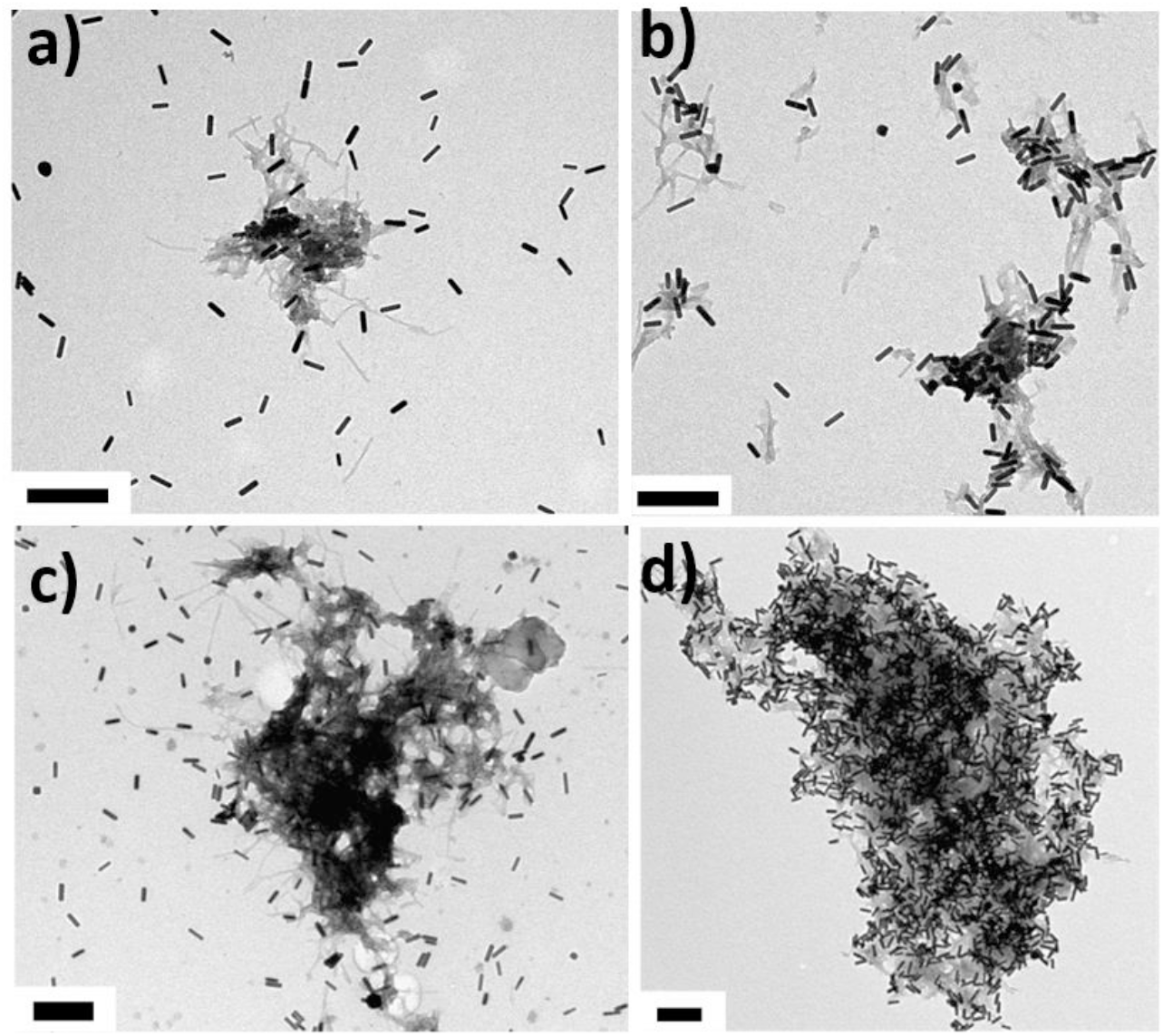
Electron microscopy confirms specific association of Abs-GNRs with Aβ seeds and fibrils. Aβ seeds incubated overnight with a) PMA-GNRs and b) Abs-GNRs. Aβ fibrils incubated overnight with PMA-GNRs c) and Abs-GNRs d). Scale bars correspond to 200 nm.

### Seeds-Abs-GNR complexes are stable and non-toxic

Seeds-Abs-GNR complexes show stability assessed by absorption spectra in mouse serum, water and cell media up to 2 days (Fig 4a and Fig S8). To test viability, SH-SY5Y cells were exposed to PMA-GNRs, Abs-GNRs, seeds-Abs-GNRs and Aβ seeds for 24 h with viability assessed using the resazurin assay (Fig 4b). Results show that the presence of 1.5 nM and 6 μM of seeds-Abs-GNRs (Fig 4b, orange) can be employed in the developed optical probe with no significant effect on cell viability. These results are in good agreement with previous studies on toxicity of gold nanorods [28, 38] and Aβ [39].

**Fig 4.**
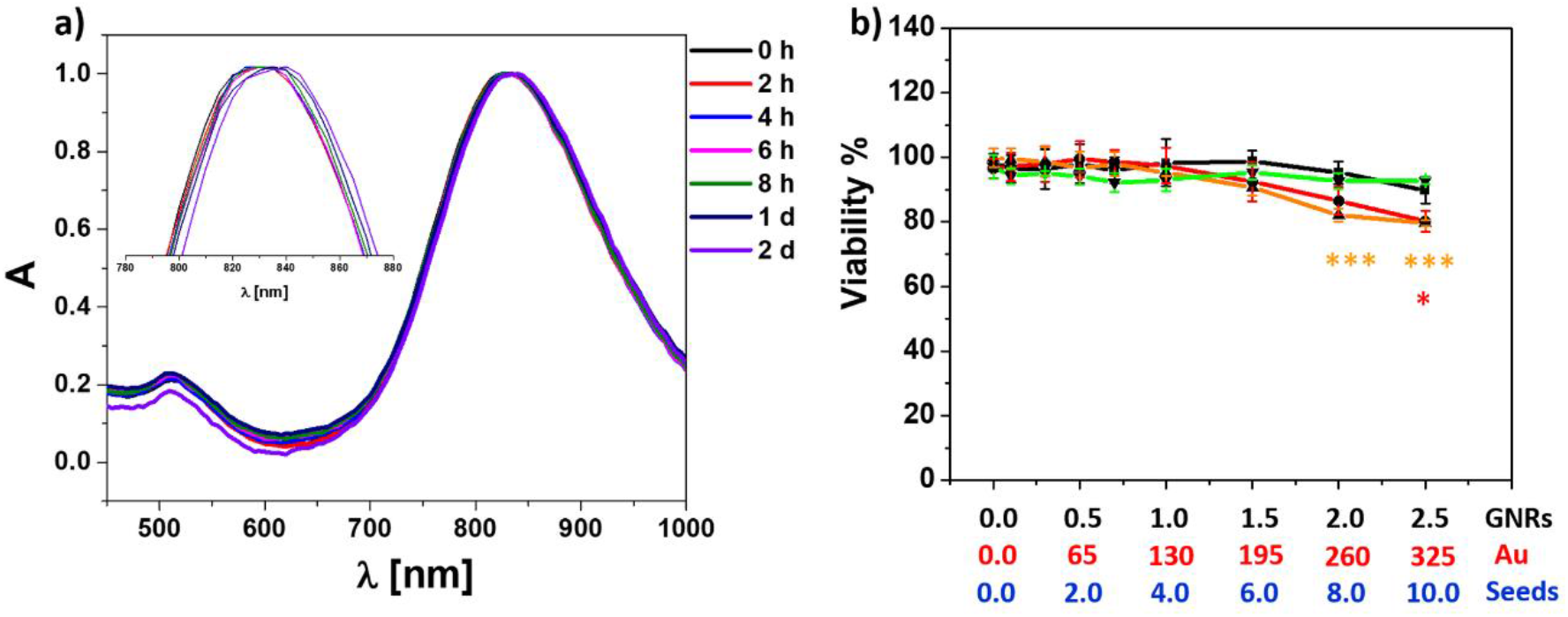
Gold nanorod complexes show stability and no toxicity under physiological conditions. (a) Normalized absorption spectra for pre-formed seeds-Aβ-GNRs in mouse serum at different time points (up to 2 d). (b) Cellular viability (%) as determined by the resazurin assay with SHSY5Y cells exposed to PMA-GNRs (black), Abs-GNRs (red), seeds-Abs-GNRs (orange) and Aβ-seeds alone (green) for 24 h at the indicated concentrations of gold nanorods (nM), gold (μg/mL) and Aβ seeds (μM). Data presented as mean ± SD from 3 wells per condition and analysed statistically using analysis of variance and Tukey’s post hoc test with p<0.05. *p<0.05, ***p<0.0001 compared to cells alone with colour corresponding to condition tested.

### Seeds-Abs-GNR-complexes retain seeding ability and can be integrated into fibrillar networks

The formation of Aβ40 fibrils is nucleation-dependent and addition of pre-formed Aβ40 seeds results in a concentration-dependent reduction in lag time (Fig 5a). Addition of seeds-Abs-GNR complexes shows a similar concentration-dependent reduction in lag time confirming they have retained the ability to seed Aβ40 fibrillation (Fig 5b). Addition of PMA-GNRs had no effect on fibril formation (Fig S9) consistent with lack of association between PMA-GNRs and Aβ40. Addition of Abs-GNRs in the absence of pre-formed seeds resulted in increased Aβ-40 fibrillation lag time and reduced fibril fluorescence intensity (Fig S8) which may be caused by depletion of Aβ from solution via binding to Abs-GNRs. TEM was used to confirm that Aβ fibrils formed in the presence or absence of seeds-Abs-GNRs show no morphological differences, with gold nanorods observed within fibrillar networks, suggesting incorporation of bound seeds during the fibrillation process (Fig 5, c and d).

**Fig 5.**
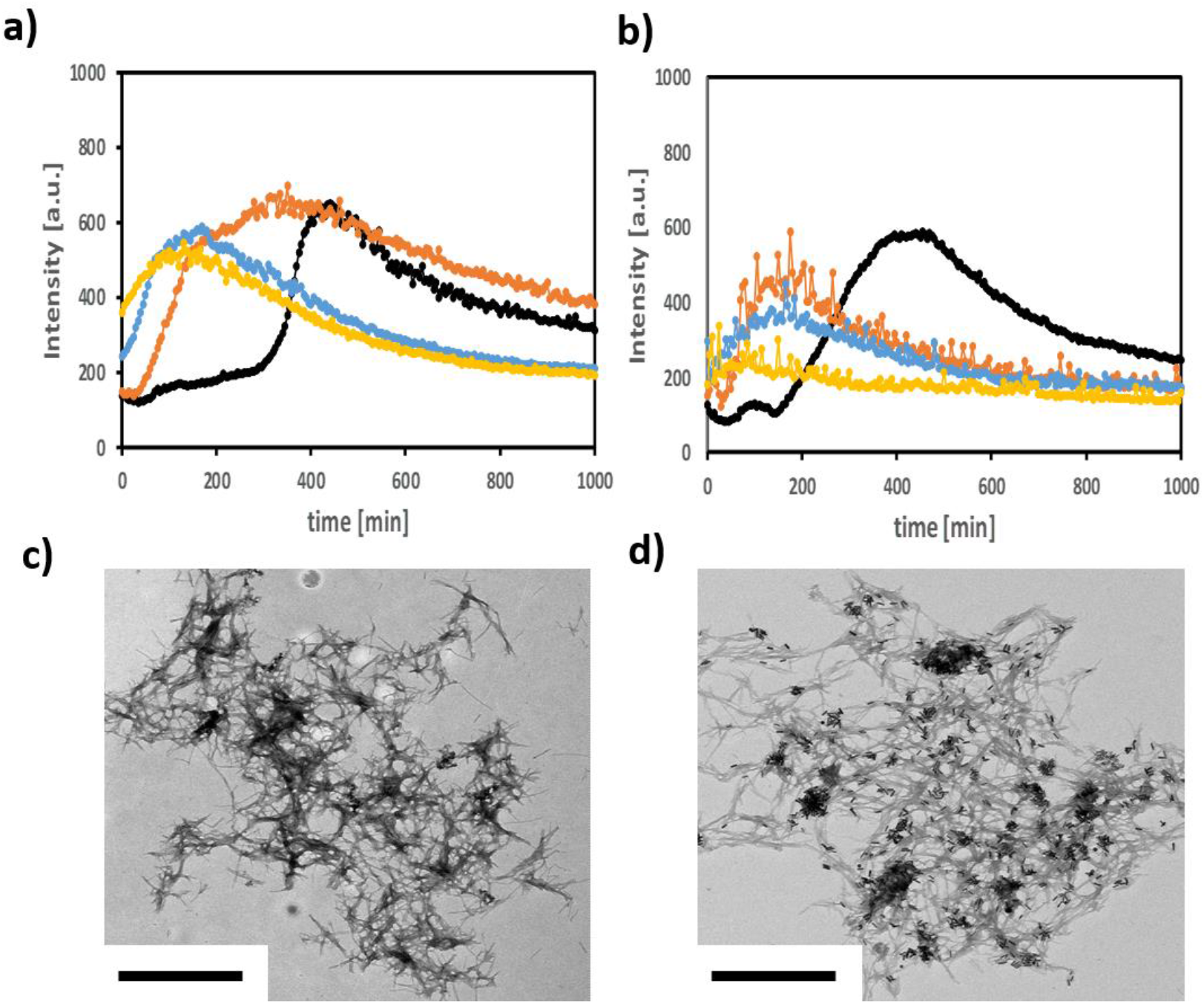
Seeds-Abs-GNRs complexes retain seeding ability and can be integrated into fibrillar networks. a) Aggregation kinetics assessed by ThT fluorescence of 10 μM Aβ40 in PBS (pH 7.4, 37°C) in absence (black) or presence of seeds at varying concentrations; 0.1 μM (orange), 0.5 μM (light blue), and 1 μM (yellow). b) ThT fluorescence of 10 μM Aβ40 incubated in absence (black) or presence of pre-formed seeds-Abs-GNRs formed from 0.4 nM of Abs-GNRs with increasing concentrations of seeds as above 0.1 μM (orange), 0.5 μM (light blue), and 1 μM (yellow). TEM images of 10 μM Aβ40 incubated in PBS, pH 7.4 for 10 h at 37 °C in the presence of c) plain seeds and d) seeds-Abs-GNRs, confirming that seeds-Abs-GNR complexes are integrated within fibrillar networks. Scale bars correspond to 1 μm.

### Optoacoustic imaging of seeds-Abs-GNR complexes

MSOT data was collected using agar phantoms to test sensitivity and optical properties of seeds-Abs-GNR complexes. A range of seeds-Abs-GNR complexes concentrations (5-500 pM) were imaged and recorded at wavelengths 680 to 950 nm. As expected, the strength of the photoacoustic signal detected by MSOT correlates with the absorbance over this range of wavelengths (Fig 6a). This is associated with a gradual change in the MOST signal intensity at different wavelengths as shown in the single wavelength images (Fig 6b). The MSOT signal was found to be linearly dependent on concentration (Fig 6c). At 5 pM, the lowest tested concentration, the unmixed MSOT signal is still detectable separately from the background noise of the agar phantom (Fig 6d) which is encouraging for future applications. Previous studies injected Aβ seeds at concentrations of 10-20 ng/μl.[10] We have made our Aβ-Abs-GNR complexes consistent with this concentration of ∼2 μM Aβ and 500 pM gold nanorods. MSOT data suggests that even with dilution of seeds upon injection that may occur sufficient signal would remain to enable tracking of seeds *in vivo*. Estimating precisely the *in vivo* limit of detection is a highly complex issue because sensitivity is affected by tissue depth, the absorption profiles of various organs, attenuation by tissues, as well as the number of wavelengths selected for acquisition [40, 41]. Further calibration work *in vivo* will be required to precisely establish the amount of Abs-GNRs that can be tracked.

**Fig 6.**
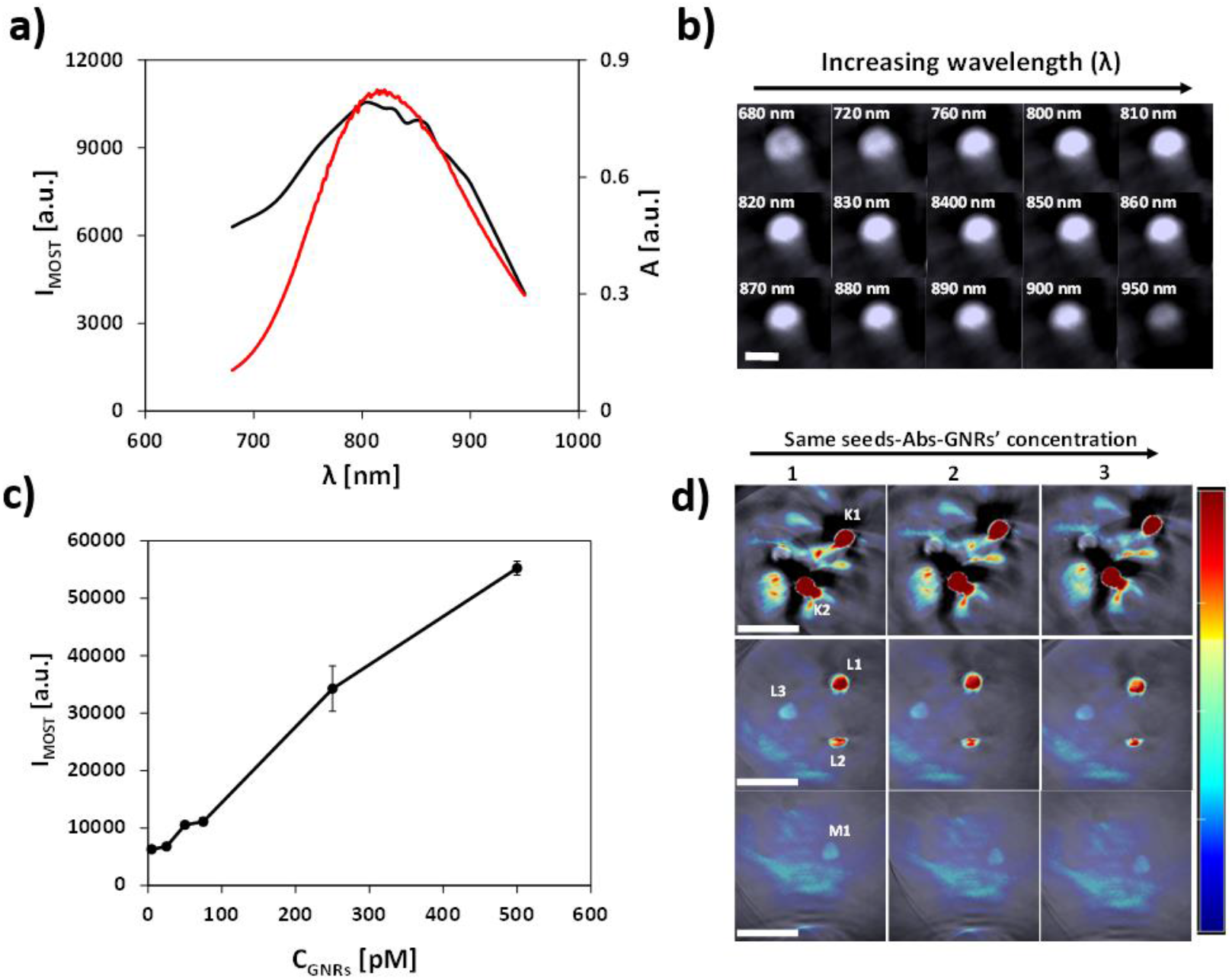
Phantom MSOT experiments. a) Comparison of MSOT intensity signals (black) and UV spectrum (red) of Aβ-seeds-GNRs agar phantom. b) Corresponding single wavelength MSOT images (680 - 950 nm). Scale bar is 3 mm for all images. c) Average MSOT signal intensities collected from 3 different positions for increasing concentrations of seeds-Abs-GNRs (5-500 pM). d) Corresponding multispectral unmixed MSOT images. Concentration of seeds-Abs-GNRs; Top-500 pM (K1), 250 pM (K2), Middle-75 pM (L1), 50 pM (L2), 25 pM (L3), Bottom-5 pM (M1). Images were reconstructed from raw signal data captured with the MSOT system at 3 different positions (1, 2 & 3) with a logarithmic colour bar. Scale bars are 10 mm. Colour scale range is 1.2 × 10^3^ to 1.2 × 10^4^ MSOT au (same scale is used in all MSOT images).

## Conclusions

In this work we have exploited the selectivity of antibodies conjugated to gold nanorods to bind to pre-formed Aβ seeds to design an optoacoustic imaging tool for monitoring the propagation of Aβ seeds *in vivo*. The resulting probes have been characterised to be stable, non-toxic and biocompatible while retaining their ability to seed fibril formation comparable with that observed by seeds alone. Preliminary MSOT data demonstrate that the optical signature of gold nanorods was preserved in the complexes and that signal could be detected at concentrations as low as 5 pM. These complexes provide a suitable optical probe to monitor the propagation of Aβ-seeds by MSOT in an *in vivo* model to contribute to a better understanding of the role of Aβ seeds in Alzheimer’s disease progression.

## Acknowledgements

We thank BioLegend for their support with providing antibodies for this project and Alzheimer’s Research UK for pilot project funding (ARUK-PPG2017B-019).

## Supporting Information

**Table S1.**
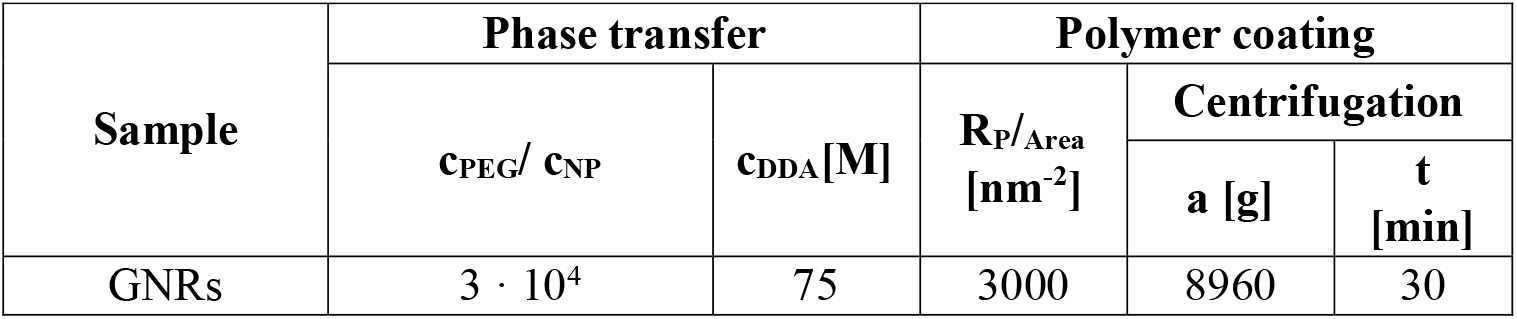
Experimental conditions used for the phase transfer, polymer coating and anti-Aß anybody conjugation. c_NP_, c_DDA_ and c_PEG_ refers to the concentration of gold nanorods, dodecylamine and PEG, respectively. R_p/Area_ refers to the number of PMA monomer added per nm^2^ of effective NP surface. a refers to the centrifugation acceleration (g = 9.81 m/s^2^) and t refers to the centrifugation time.

**Table S2.** EDC/sulfo-NHS coupling reaction of Abs-Aβ to PMA-GNRs under different concentrations of EDC/sulfo-NHS at a fixed concentration of Abs-Aβ (sampled coded “a”) or different concentrations of Abs-Aβ at a fixed concentration of coupling reagents (samples coded “b”). The ratio between EDC and sulfo-NHS was always kept constant at 1:2, respectively.

| **Condition** | **Abs-Aβ**<br>**[μg/ mL]** | **EDC**<br>**[mmoles]** | **sulfo-NHS**<br>**[mmoles]** | **LSPR shifting**<br>**[nm]** |
| --- | --- | --- | --- | --- |
| 1a | 0 | 0 | 0 | Control |
| 2a | 50 | 0 | 0 | 0 |
| 3a |  | 0.5 | 1 | ~2 |
| 4a |  | 1 | 2 | ~3 |
| **5a** |  | **2** | **4** | **~5** |
| 6a |  | 4 | 8 | ~5 |
| 1b | 0 | 0 | 0 | Control |
| 2b | 5 | 2 | 4 | 0 |
| 3b | 20 |  |  | ~2 |
| **4b** | **50** |  |  | **~5** |
| 5b | 75 |  |  | ~5 |

**Table S3.** Hydrodynamic diameter d_h_ [nm] in number (d_h(N)_) and intensity (d_h(I)_), Z-average (d_h(Z)_), polydispersity index (PDI) and ζ -potential values (ζ) for gold nanorods along their surface modification and upon conjugation with Aβ seeds. Size values for gold nanorods, although without any physical meaning because they are not spherical NPs, are included to show that the colloidal stability was not compromised in the different steps of the polymer coating.

| Sample | Coating | $d_{h,N}$ [nm] | $d_{h,Z}$ [nm] | PDI | $d_{h,I1}$ [nm] | $d_{h,I2}$ [nm] | $\zeta$ [mV] |
| --- | --- | --- | --- | --- | --- | --- | --- |
| **GNRs** | CTAB | $3.65 \pm 0.05$ | $7.87 \pm 0.25$ | $0.3 \pm 0.01$ | $6.62 \pm 0.01$ | $99.49 \pm 1.65$ | $+34.1 \pm 1.15$ |
| | DDA | $2.75 \pm 0.03$ | $8.0 \pm 0.19$ | $0.3 \pm 0.01$ | $4.77 \pm 0.02$ | $79.82 \pm 1.06$ | - |
| | PMA | $3.90 \pm 0.1$ | $10.38 \pm 0.1$ | $0.4 \pm 0.008$ | $7.69 \pm 0.16$ | $89.54 \pm 1.87$ | $-28.8 \pm 0.26$ |
| | Abs | $4.5 \pm 0.3$ | $12.29 \pm 0.09$ | $0.4 \pm 0.006$ | $8.25 \pm 0.2$ | $97.54 \pm 1.2$ | $-17.7 \pm 1.75$ |
| **Plain Aβ Seeds** | - | - | - | - | - | - | $-44.2 \pm 1.10$ |
| **Aβ Seeds-GNRs** | - | - | - | - | - | - | $-43.7 \pm 0.30$ |

**Figure S1.**
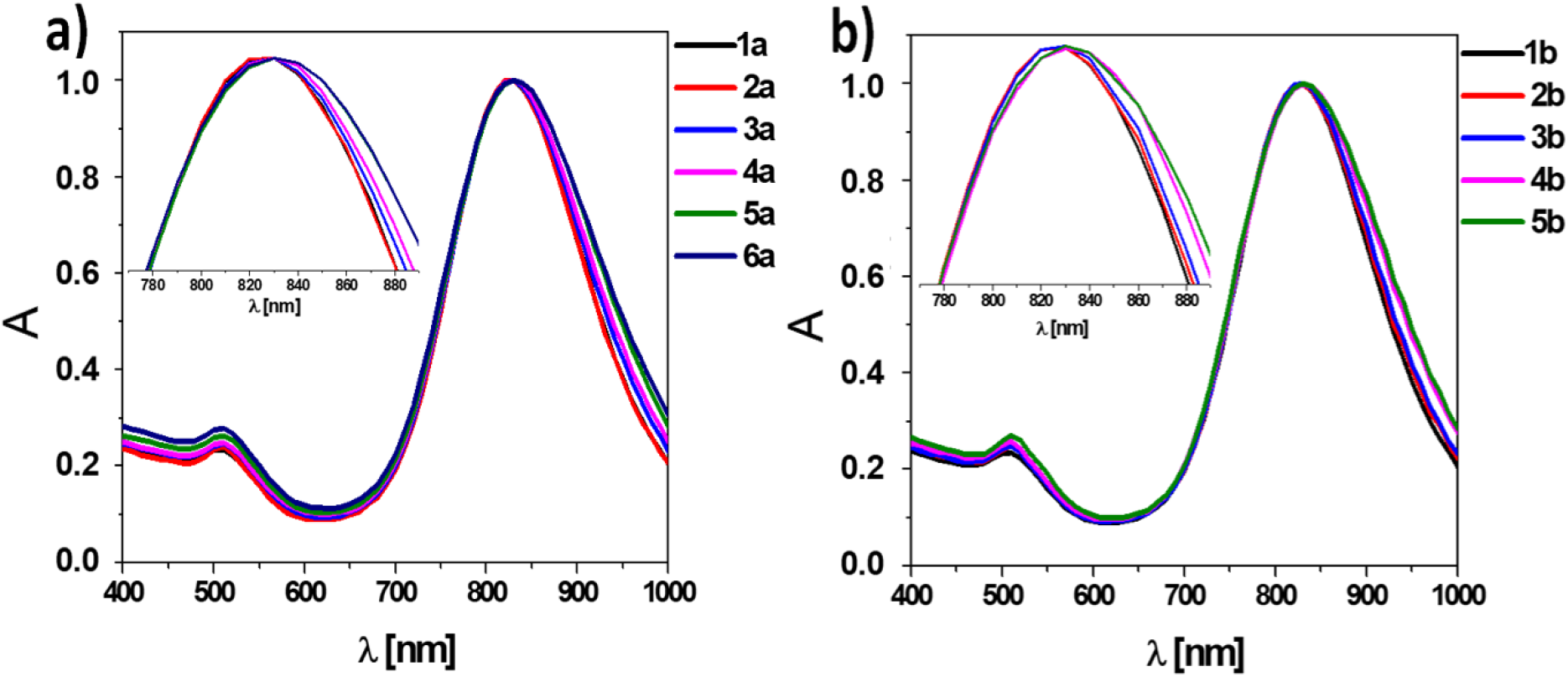
Normalized absorption spectra for PMA-GNRs before and after conjugation with Abs-Aβ under different experimental conditions using EDC/sulfo-NHS. a) The concentration of EDC/sulfo-NHS was varied while the concentration of Abs-Aβ was fixed. b) The concentration of Abs-Aβ was varied while the concentration of EDC/sulfo-NHS was kept constant. The experimental conditions are listed in Table S2.

**Figure S2.**
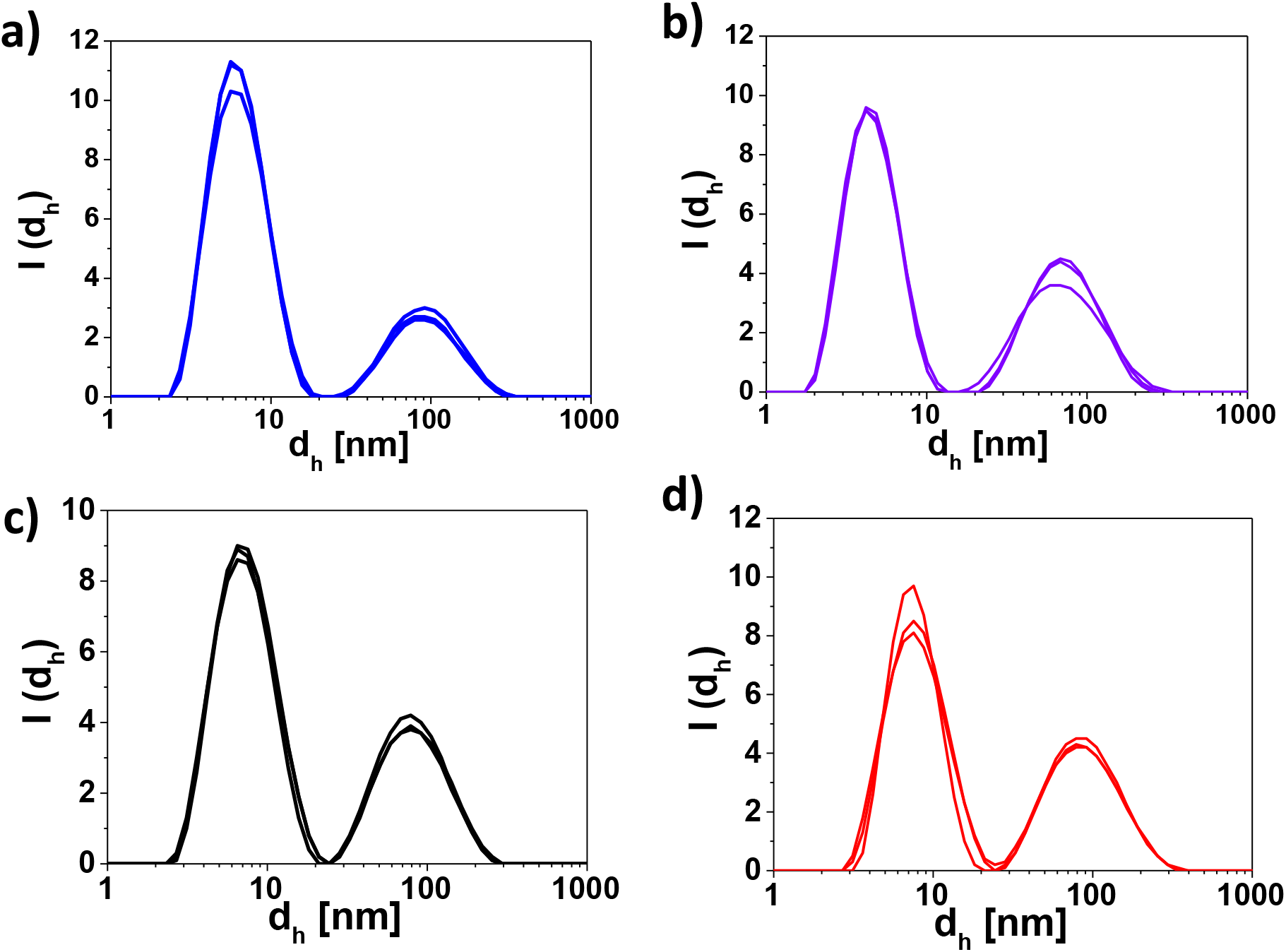
Intensity distribution of hydrodynamic diameters (I(d_h_)) for gold nanorods along their surface modification. a) CTAB-capped; b) DDA-capped; c) PMA-coated and d) after conjugation with anti-Aβ antibody.

**Figure S3.**
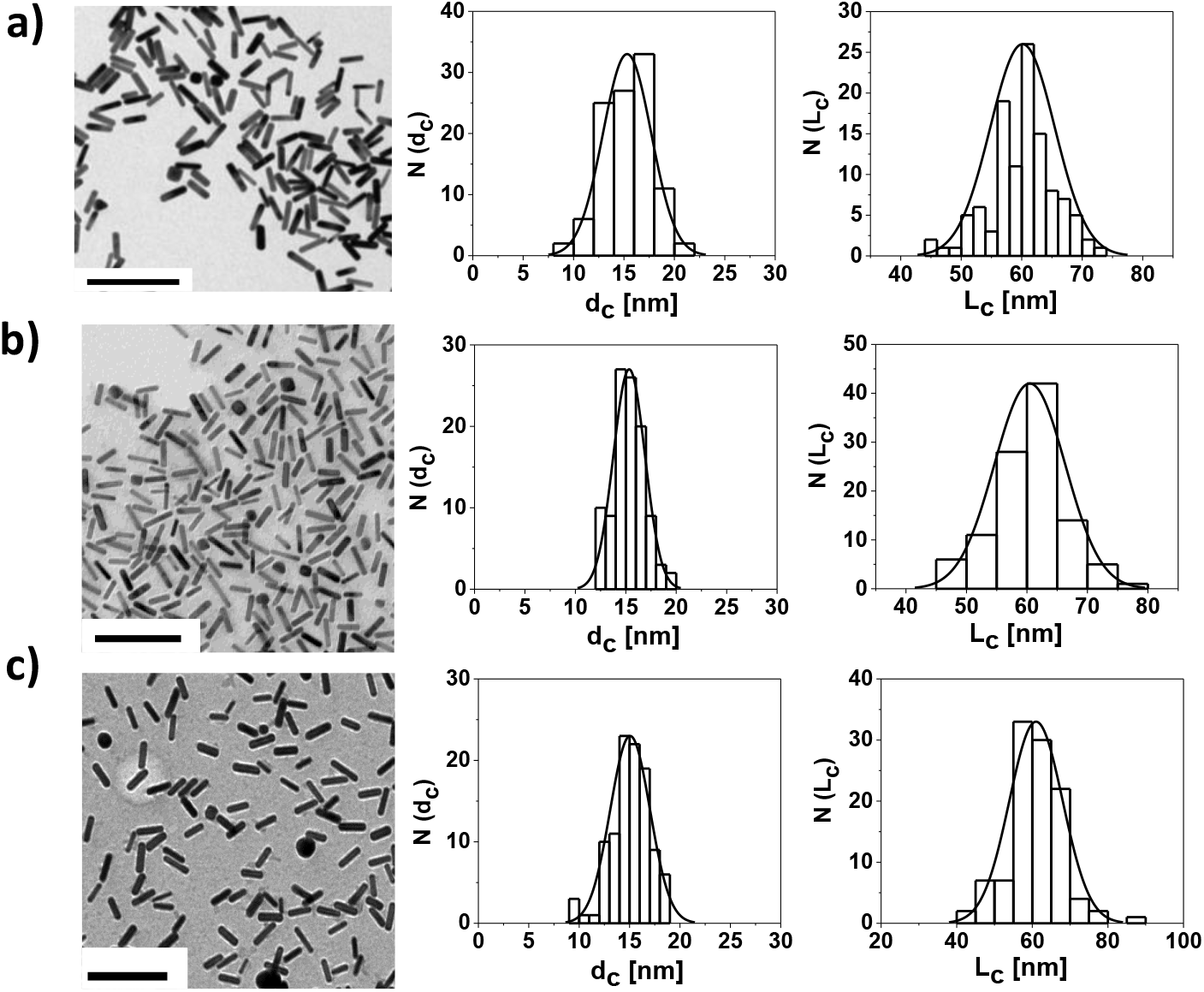
TEM images of gold nanorods and their corresponding size distribution histograms after different steps of surface modifications, plotted as number of NPs. N(d_c_) that have a core diameter of d_c_ (histograms to the left) and as number of NPs N(L_c_) that have a core length of L_c_ (histograms to the right). A) CTAB coated gold nanorods with d_c_ = 15.31 ± 2.39 nm, and L_c_ = 60.13 ± 5.33 nm. B) PMA coated gold nanorods with d_c_ = 15.25 ± 1.68 nm, and L_c_ = 59.90 ± 5.85 nm. C) Ab conjugated gold nanorods with d_c_ = 14.91 ± 1.97 nm, and L_c_ = 60.987.06 nm. The scale bars correspond to 200 nm.

**Figure S4.**
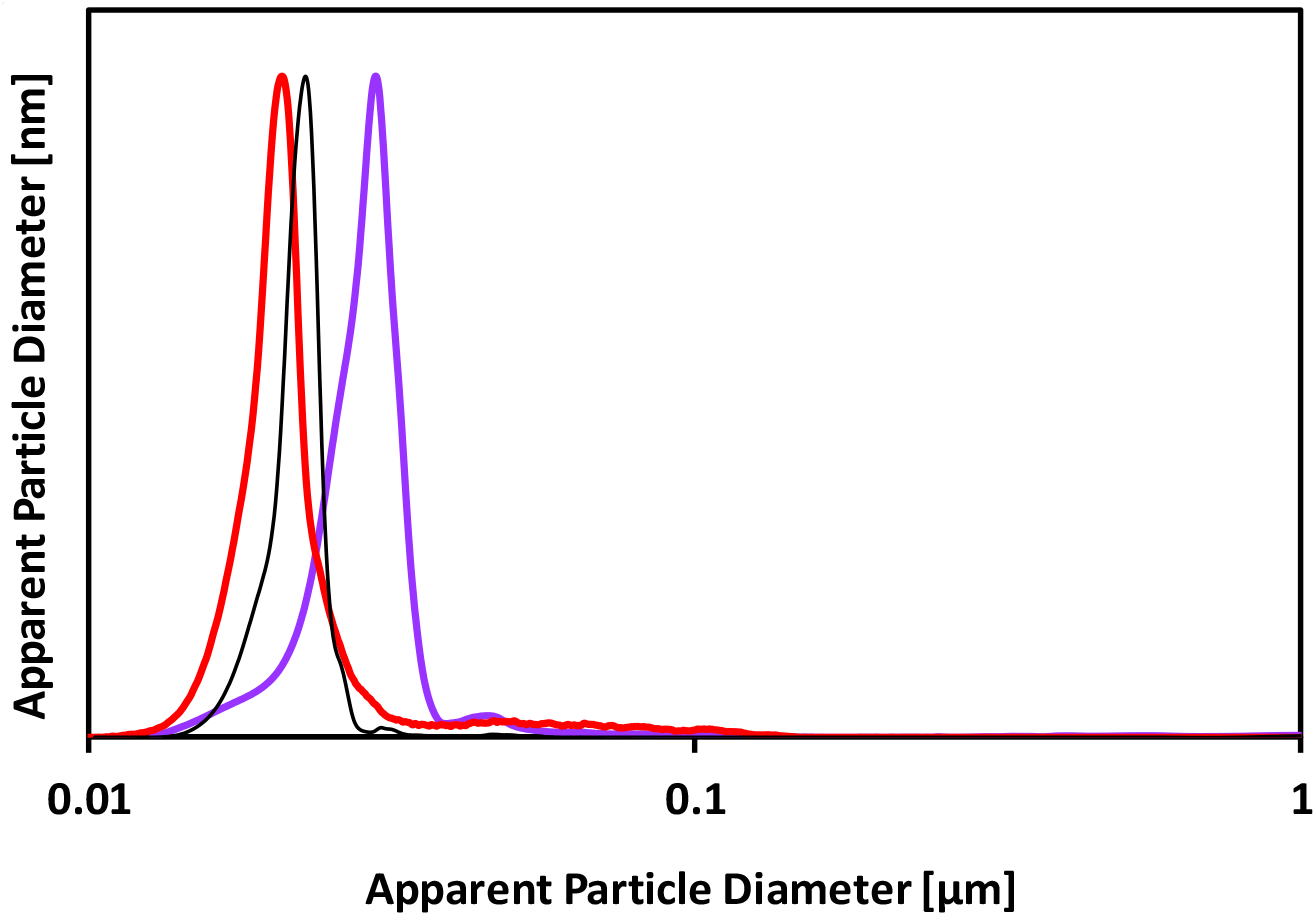
Apparent size distributions measured by analytical disc centrifugation (CPS). CTAB-capped gold nanorods (purple), PMA-coated GNRs (black) and Aβ antibodies conjugated PMA-GNRs (red). The average values are 29.8 ± 0.2 nm, 22.79 ± 0.12 nm and 20.81 ± 0.18 nm respectively.

**Figure S5.**
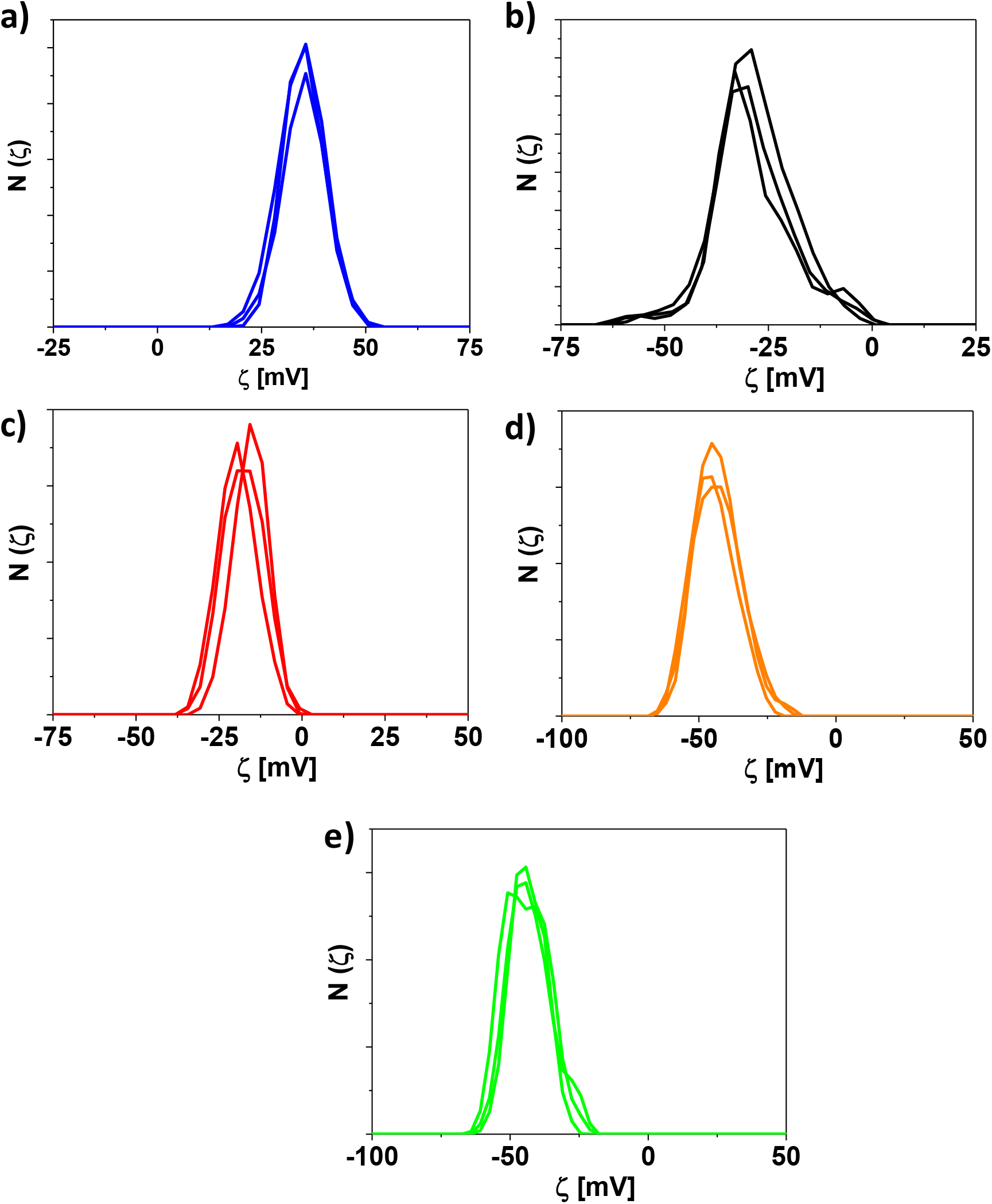
ζ-potential distribution (N(ζ)) results during gold nanorods surface modification process. a) CTAB-capped; b) PMA-coated; c) after conjugation with anti-Aβ antibodies and d) after complex formation of Abs-GNRs attached to Aβ seeds, e) Aβ seeds alone.

**Figure S6.**
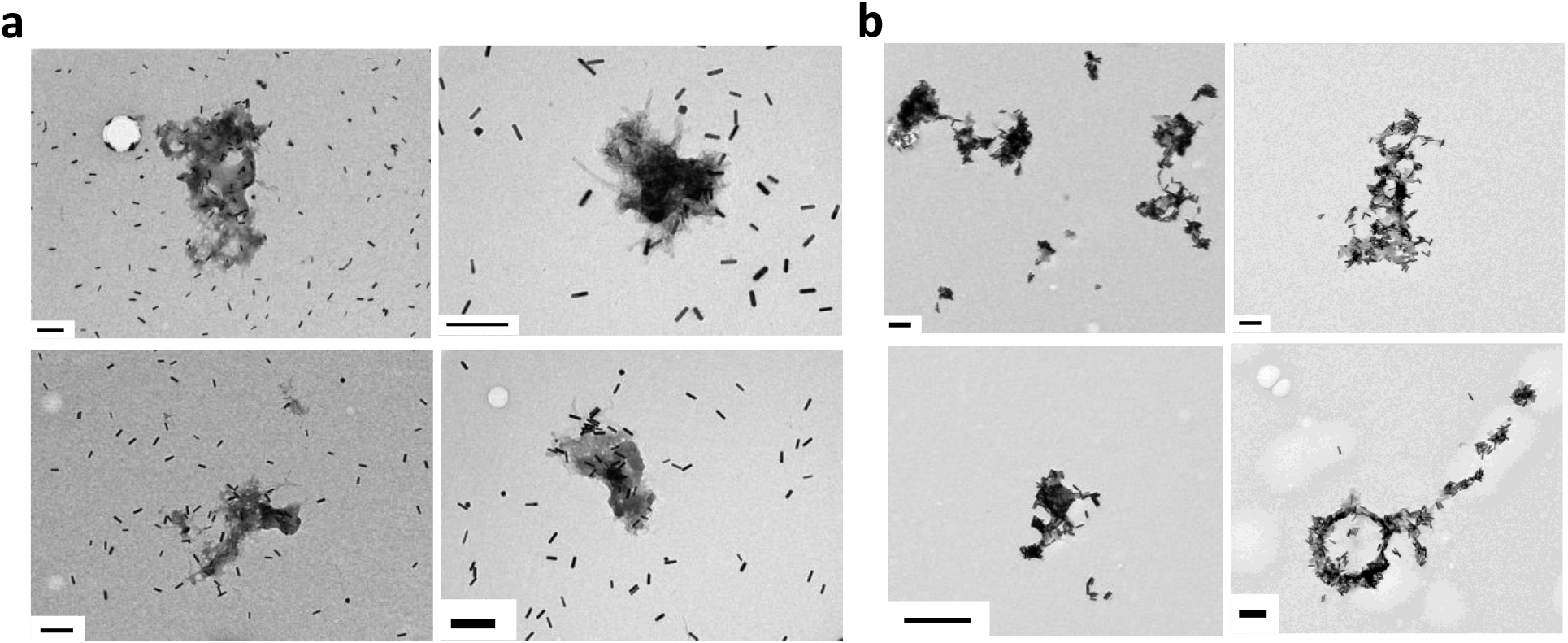
Additional TEM images. a) PMA-GNRs and b) Abs-GNRs after incubation with Aβ seeds overnight. The scale bars correspond to 200 nm.

**Figure S7.**
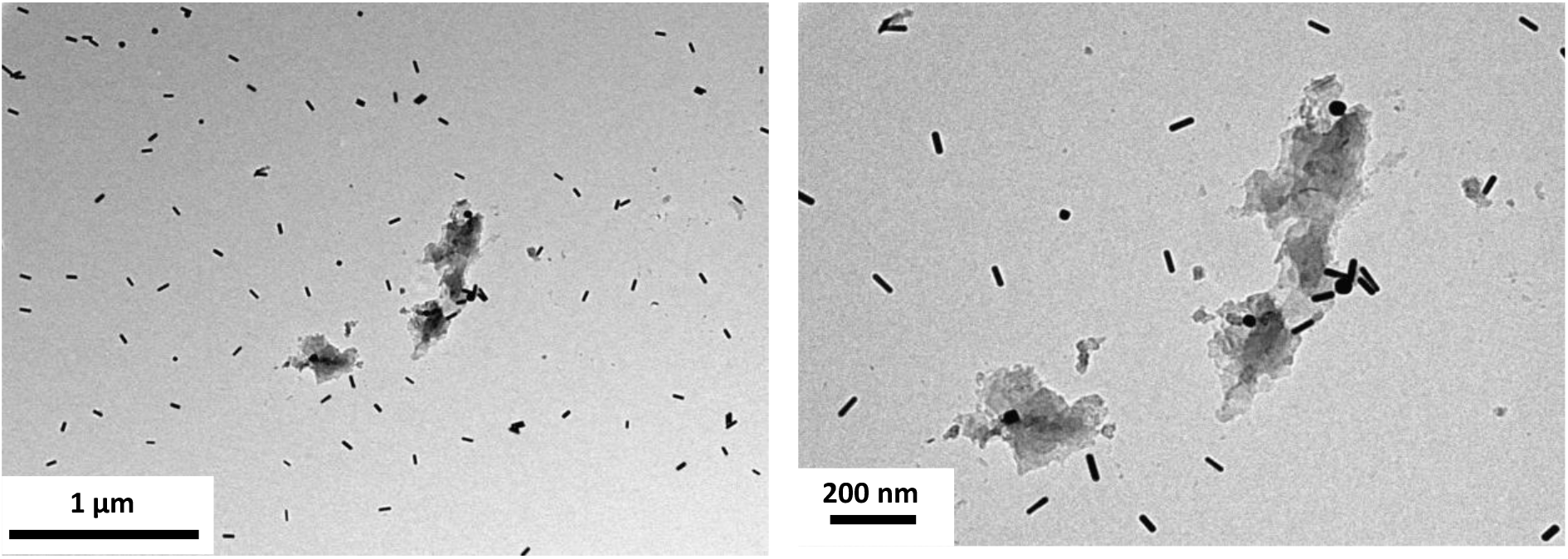
TEM images of Abs-GNRS incubated overnight with seeds formed from a control amyloidogenic protein medin. Scale bars are as shown.

**Figure S8.**
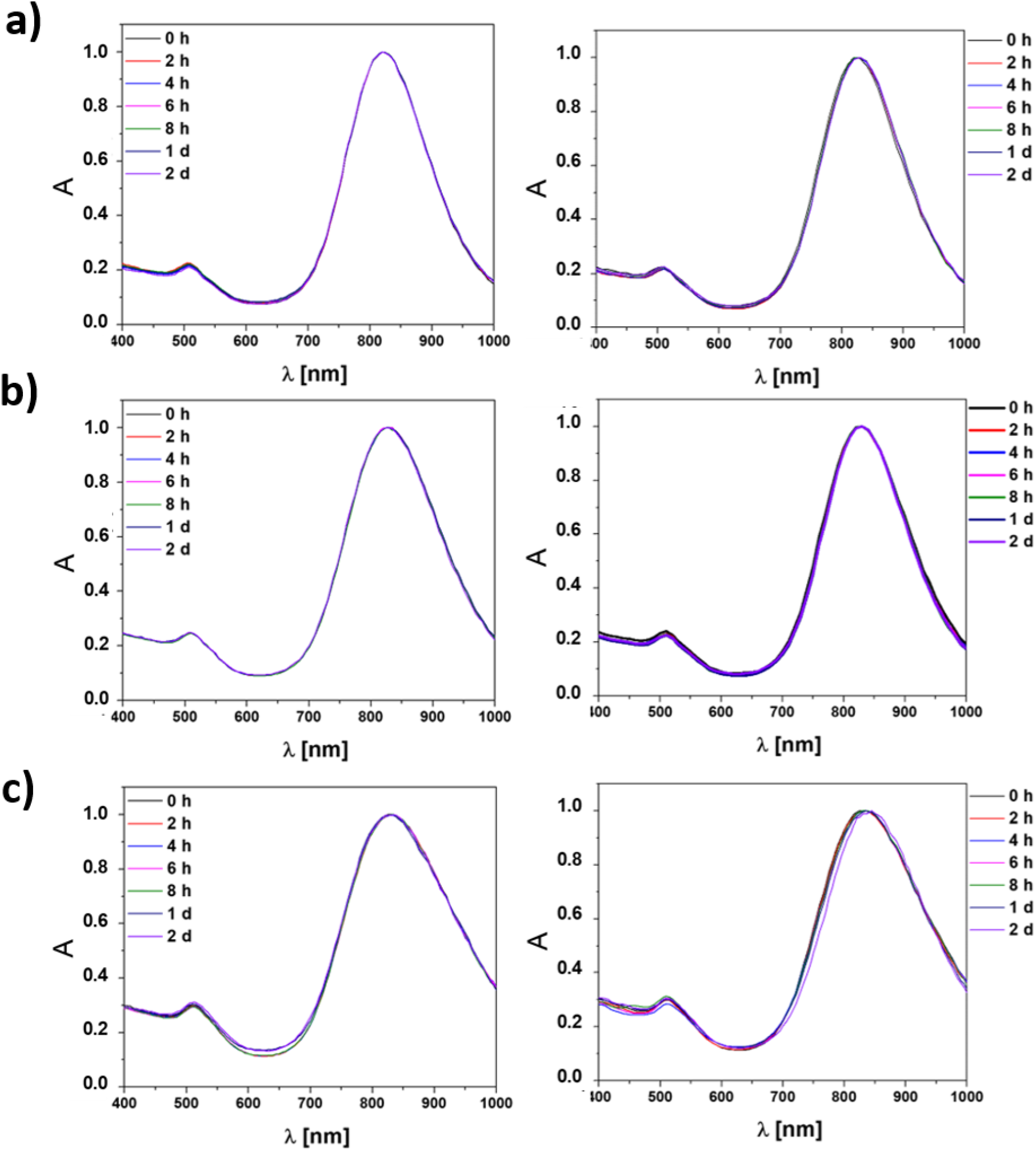
Normalized absorption spectra. a) PMA-GNRs, b) Abs-GNRs, and c) seeds-Abs-GNRs in water (left) and cell media (right) at different point of times (up to 2 days).

**Figure S9.**
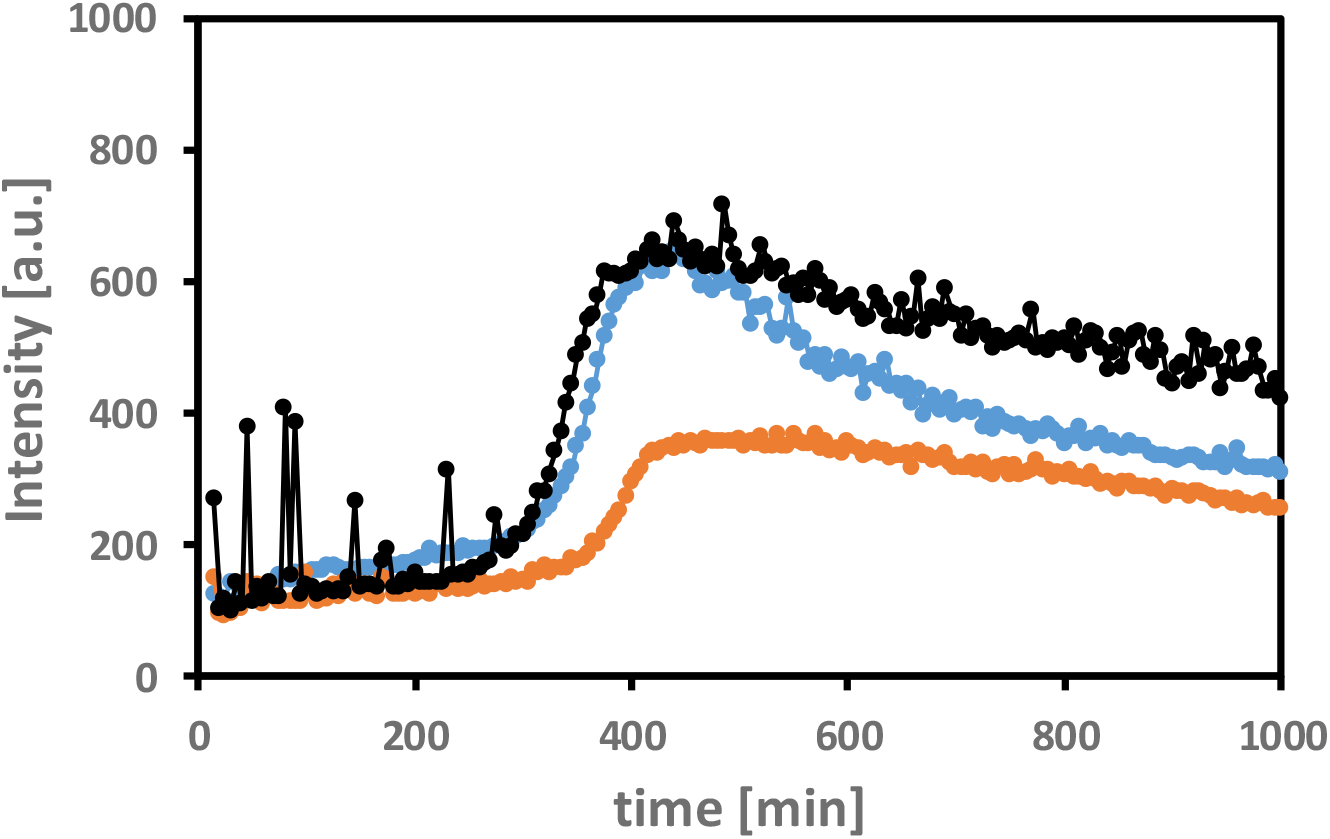
PMA-GNRs do not bind to or alter Aβ40 fibrillation. ThT fluorescence of 10 μM Aβ40 in PBS (pH 7.4, 37°C) alone (black), and in the presence of PMA-GNRs (light blue) or Abs-GNRs (orange).

## Appendix

### I. Calculating the amount of PMA needed for phase transfer of NPs from organic solvent to aqueous medium

To determine the amount of polymer which is needed for the coating of the NPs, a calculation based on the total NP surface to be coated was done following the calculations previously reported by Pellegrino et al.,^i^in which the total effective surface area of one nanoparticle (A_eff_) is calculated in agreement with its shape. Nanorodswere considered as cylinders to simplify the calculations. In this case, the effective diameter (d_eff_) value included the diameter of the Au core as determined by transmission electron micoscopy (TEM) (d_c_, cf. section 3) plus two times the assumed thickness of the surfactant shell (L_s_ = 1 nm). Once the surface of one NP is determined then, knowing the NPs concentration (c_NP_) and the volume of the NP solution to be coated (V_NP,sol_), the total area (A_tot_) of the NPs in the solution can be determined by the formula (S1):

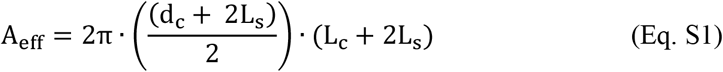

A_eff_ is the effective surface area of one NP including the organic surfactant layer, d_c_ is the diameter of the Au core and L_c_ is the length of Au core of the gold nanorodsas determined by TEM. L_s_ corresponds to the thickness of the surfactant layer and d_eff_is the effective diameter, equal to the NP core diameter determined by TEM (d_c_) plus two times the assumed thickness of the surfactant shell (L_s_ = 1 nm). The total effective surface area (A_tot_) of NPs thus is:

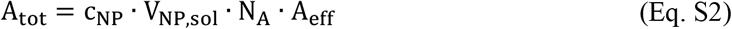

c_NP_ is the NP [M] concentration of the NP solution used of the coating. V_NP,sol_ [L] refers to the volume of the NP solution, and N_A_ is the Avogadro constant. Then, the amount of polymer needed to be added to the NP solution can be determined by formula S3.

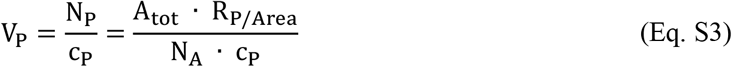

N_P_ [mol] is the amount of polymer molecules (given in terms of polymer monomers) needed to coat all the NPs in solution. cP [M] is the concentration of the polymer stock solution. VP [L] is the volume of polymer stock solution which needs to be added to the NP solution. A_tot_ [nm^2^] is the total surface area of all the colloidal NPs in the solution and R_P/Area_ is the number of monomers to be added per nm^2^of effective NP surface. N_A_ is the Avogadro constant.

### II. Conversion of gold nanorods’ molar concentration to mass concentration

To convert the molar values of gold nanorods determined by UV/vis spectroscopy (cf.section 2.2, Eq. 1) into mass concentrations [μg/mL], the molecular weight for each NPs was calculated based on the volume equation of a rod (Eq. S4).

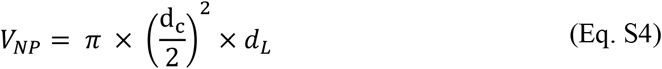

The nanorods molecular weight (M_w_) was calculated by following Formula S5:

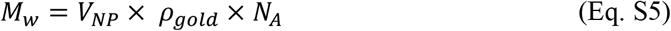

Where p is the gold density and NAis the Avogadro number.

With neglecting the contribution of the polymer coating to the mass of the NPs,^ii^The conversion of mol to mass can be caculated by using the following Formula S6:

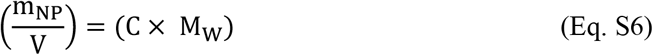

Where C is the concentration [M], m is the mass [mg], M_w_ is the molecular weight [g/mol] and V, the volume of the NPs solution [mL].

